# Summertime Fog and Dew Impacts on Vegetation Depend on Aridity

**DOI:** 10.64898/2026.07.21.739924

**Authors:** Ioannis Lolos, John T. Abatzoglou, Tyson J. Terry

## Abstract

Rainfall and vapor pressure deficit (VPD) are well-studied hydrological variables that largely determine aboveground net primary production (ANPP) in most ecosystems. Meanwhile, the impacts of another important part of the hydrologic cycle, non-rainfall water from fog and dew, remain poorly understood at the ecosystem level. To fill this gap, we used meteorological variables measured at weather stations along with satellite-derived vegetation greenness data from surrounding areas to examine how fog and dew frequency affect summer plant growth across the contiguous United States. Our analysis shows that, even after accounting for precipitation, VPD, and land-cover type, fog and, more so, dew enhanced vegetation productivity in water-limited regions. In contrast, non-rainfall water had a neutral or negative impact on plant growth in humid regions, with fog showing the strongest and most widespread negative effects. Taken together, our findings reveal that summertime non-rainfall water has differential effects on vegetation that are largely determined by ecosystem-level water availability. These aridity-dependent effects of fog and dew should be considered in future ecological and agricultural studies and in assessments of projected climate impacts on vegetation.

## 1. Introduction

Non-rainfall water (NRW) represents one of the least understood hydrological components in most ecosystems (1). However, in recent years fog and dew have attracted increasing interest from the scientific community. This interest has arisen largely due to accumulating evidence that NRW benefits biota in arid and semi-arid regions (1–4), although how NRW influences vegetation productivity at a large scale and across the spectrum of water-availability regimes remains poorly understood (5).

Both fog and dew contribute to the water budget and alter plant functions in unconventional ways. For instance, a considerable fraction of the dew and fog water is deposited on aboveground plant tissues rather than directly into the soil (2, 3). This water can then be absorbed through leaves, directly increasing turgor pressure through a phenomenon known as foliar water uptake, which has been shown to occur in more than 90% of all plant species tested (6). Such leaf-wetting events have also been shown to alter photosynthesis and transpiration in various ways, depending on their timing, water availability in a system, and the specific plant species (6). Moreover, leaf-wetting can transiently decrease leaf temperatures, encourage growth of beneficial endophytes or detrimental plant pathogens, and contribute to nutrient leaching and nutrient or atmospheric pollutant deposition (3, 6). Additionally, fog alters the path, intensity, and quality of light before it reaches the canopy, which can either enhance or suppress photosynthesis and light-use efficiency depending on scale, canopy structure, leaf traits, and abundance of various resources (7–10).

Even though these insights are largely derived from localized, species-specific, and therefore hard-to-generalize studies, it seems that NRW has the potential to differentially impact vegetation. What then determines the actual net effect, and might it differ from one ecosystem to another? We hypothesize that, although many factors may modulate the net effect of NRW, ecosystem-level water balance is the primary determinant of plant responses at broad spatial scales. We expect this to be the case because, in humid ecosystems, water is not the limiting factor and leaves are wet for prolonged periods of time, making NRW redundant and in fact potentially harmful (due nutrient leaching, moisture-dependent diseases, light limitation, etc.). On the contrary, in water-limited ecosystems, NRW is expected to provide an additional source of moisture and transiently alleviate heat and evaporative stress, making it beneficial to species that are adapted to fog and/or dew.

Motivated by the above hypothesis, we conducted the first large-scale, cross-ecoregional empirical assessment of the effects of summertime fog and dew on plant growth in the contiguous United States using remotely sensed vegetation data and *in situ* meteorological observations. Although we expected aridity to be the primary determinant of plant responses to NRW, we also evaluated whether its influence varies by land-cover type, particularly because different land-cover types contain different plant communities with some more strongly shaped by anthropogenic management than others.

## 2. Results

Our final dataset included 1,080 weather stations providing broad spatial coverage across the contiguous U.S., each one with summer data over 2012–2023. The distribution of these stations, along with their fog and dew frequencies, is presented in *Fig. 1*. Both average summer fog and dew frequency showed a clear east-west spatial gradient. That is, fog occurred frequently during the summer in many parts of the eastern U.S., whereas fog almost never occurred in the western U.S. except in some coastal locations (*Fig. 1*). Dew likewise occurred much more frequently during the summer in the eastern U.S. and along the West Coast, although it also occurred in smaller amounts inland in the West (*Fig. 1*). Overall, most locations had more hours with dew than fog.

**Fig. 1.**
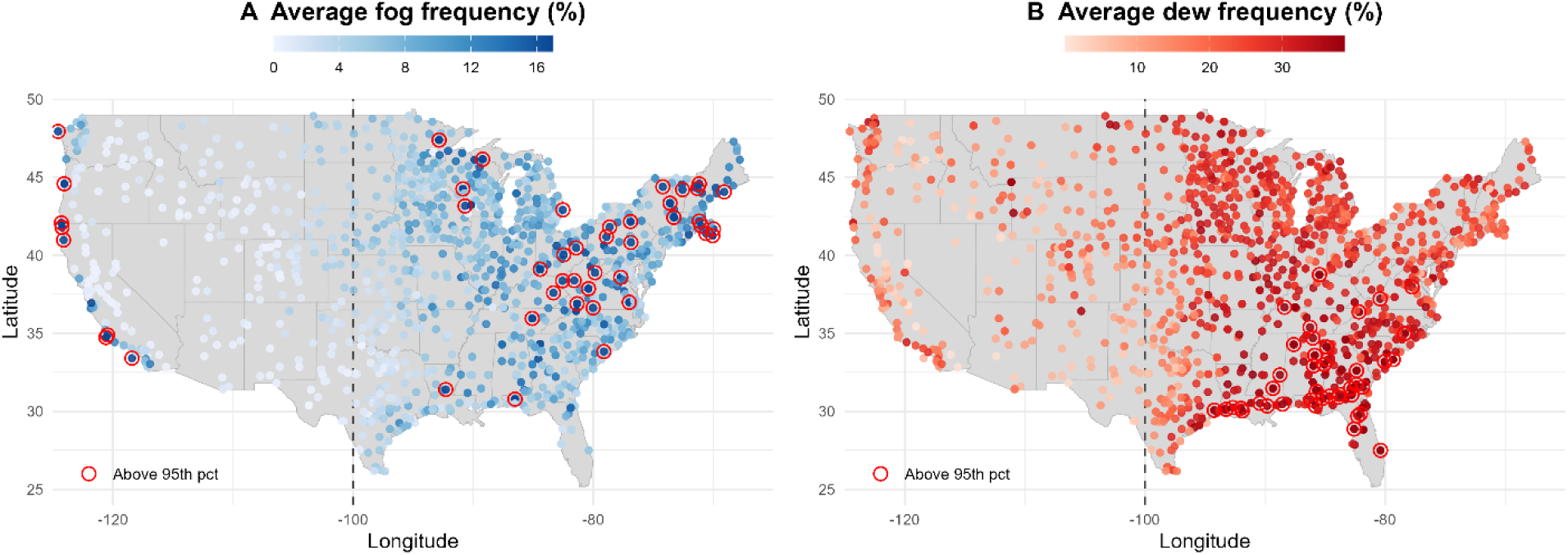
Summer fog and dew frequency for the 1,080 ASOS stations. Metrics represent averages over all summer 16-day periods during June–August, 2012–2023. Fog frequency for a given period represents the percentage of hours within that period during which fog or mist was recorded at the station. Dew frequency for a given period represents the percentage of hours within that period during which dew was estimated to occur using the Beysens equation (11).

After dividing stations into 30 equal-sized bins comprised of locations with similar summer Aridity Index (AI), for each bin separately we fit a linear mixed-effects model predicting next-period Enhanced Vegetation Index (EVI; a proxy for aboveground vegetation productivity) from prior-period EVI, current-period mean vapor pressure deficit (VPD), precipitation amount, dew frequency, fog frequency, and average cloud coverage, with a station-level random intercept. Model performance was acceptable across all 30 aridity bins (*Table S1*). Marginal R² values ranged from 0.04 to 0.43, indicating that fixed effects explained between 4% and 43% of the variance in subsequent EVI. Conditional R² values were consistently higher (0.48–0.70), reflecting additional variance explained by station-level random effects.

Precipitation and VPD determined a large portion of the EVI-derived aboveground net primary production (ANPP) during the summer, but in some regions more so than in others (*Fig. 2)*. Specifically, 16-day precipitation amount was most beneficial to plant growth in semi-arid and mesic climates, with its effects diminishing in the humid U.S. regions of the East and the hyper-arid regions of the West (where virtually no precipitation occurs during the summer). Mean VPD exerted a negative influence on plant growth in mesic climates, with a lesser but still statistically significant effect in humid regions and a negligible effect in the arid and some semi-arid locations of the West.

**Fig. 2.**
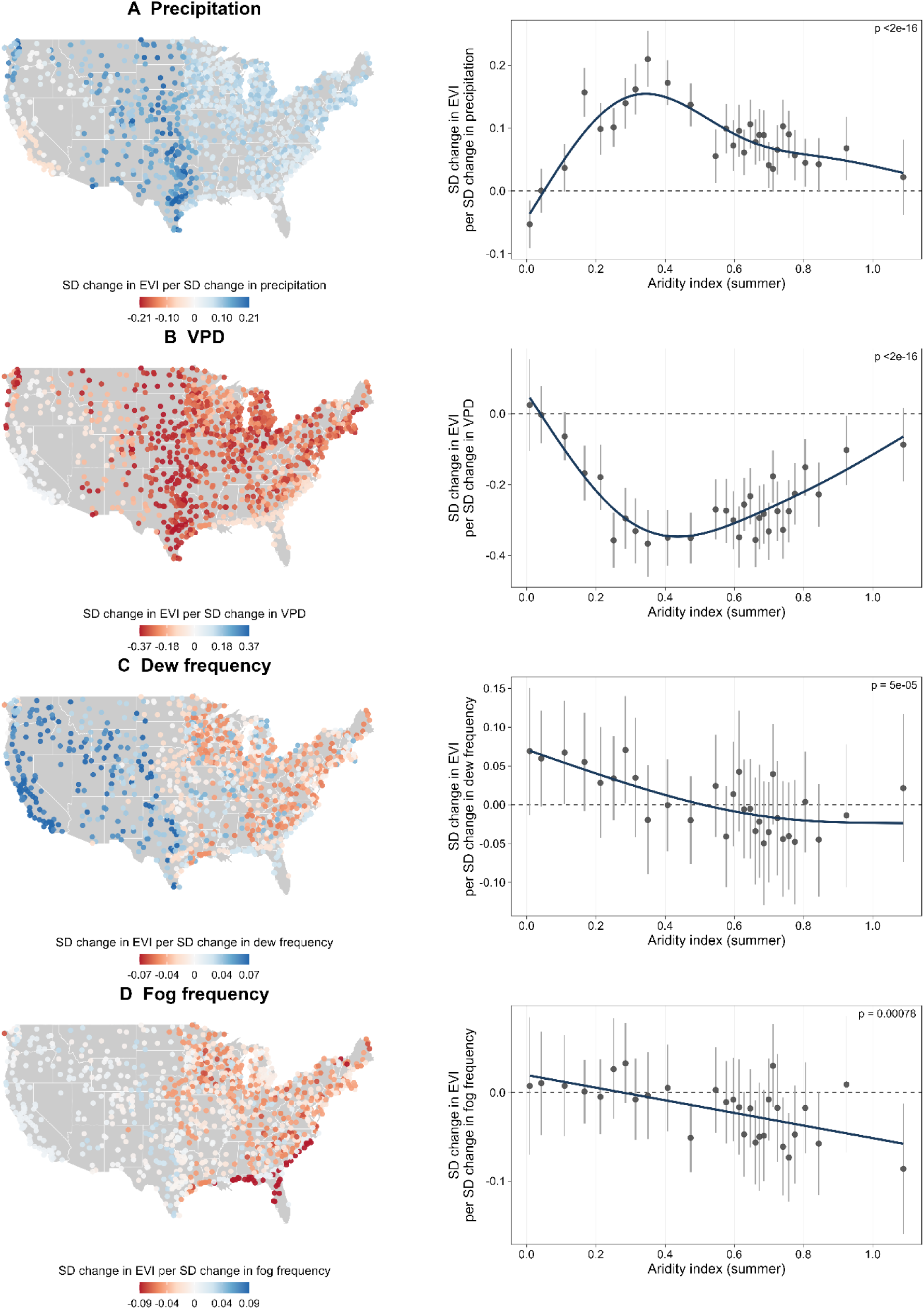
Vegetation responses to precipitation (A), VPD (B), dew frequency (C), and fog frequency (D) across the contiguous U.S. and along the aridity gradient. For each predictor, the left panel shows the spatial distribution of linear mixed-effects model coefficients per station, colored by the estimate for the aridity bin each station belongs to. The right panel shows how the coefficient varies along the summer aridity index gradient across 30 bins (lower AI values indicate more arid conditions). The dark line is a generalized additive model (GAM) smooth weighted by inverse variance; the p-value in the upper right of each panel tests whether this smooth differs significantly from a flat line at zero (i.e., whether the coefficient varies systematically with aridity).

Dew frequency was a weaker predictor of next-period EVI than precipitation or VPD, yet its effect varied systematically with aridity (*Fig. 2)*. While dew displayed slightly negative to no effect on vegetation growth in the humid regions of the East, its effect became increasingly positive as summer aridity increased in the West. Fog frequency likewise displayed aridity-dependent effects on plants, with a more pronounced negative impact on vegetation growth (compared to dew) in the humid stations that became mostly neutral as aridity increased.

We split stations into ecoregions defined by land-cover type and broad geographic region (east or west of the 100th meridian) to examine whether the observed aridity-dependent effects of fog and dew on plant growth persisted after accounting for land-cover type. Of the 1,080 stations, 856 met the inclusion criteria (consistent land-cover type across all years), spanning 9 ecoregions across the contiguous U.S. (*Fig. S1*).

For each ecoregion, we fit a linear mixed-effects model predicting next-period EVI from prior-period EVI, current-period mean VPD, precipitation, fog frequency, and dew frequency, with a station-level random intercept. Model performance varied across ecoregions but was acceptable in all cases (*Table S2*). Marginal R^2^ values ranged from 0.04 to 0.51, indicating that fixed effects explained between 4% and 51% of the variance in subsequent EVI, with the lowest explanatory power in East Woody Savannas and the highest in West Open Shrublands. Conditional R^2^ values were consistently higher in all ecoregions (0.44 – 0.63), reflecting additional variance explained by station-level random effects.

Precipitation and VPD had the strongest effects on vegetation out of the hydrological variables modeled (*Fig. 3*). Specifically, increases in 16-day mean VPD were found to strongly negatively affect vegetation growth more so in humid (eastern) than water-limited (western) ecoregions, with a particularly pronounced effect in East Grasslands and East Croplands (the two ecoregions found in the middle of the summer aridity spectrum but that also have the most responsive plant communities). West Open Shrublands and West Croplands were the only ecoregions where VPD coefficients, although negative, had confidence intervals overlapping zero. This was likely related to the models’ inability to properly distinguish between the effects of VPD and dew on vegetation growth, as indicated by the high instability of the VPD coefficient when dew frequency was added to the model (*Fig. S2)*. We believe that this instability stems, in large part, from the low number of stations in these two ecoregions (11 for West Shrublands and 16 for West Croplands).

**Fig. 3.**
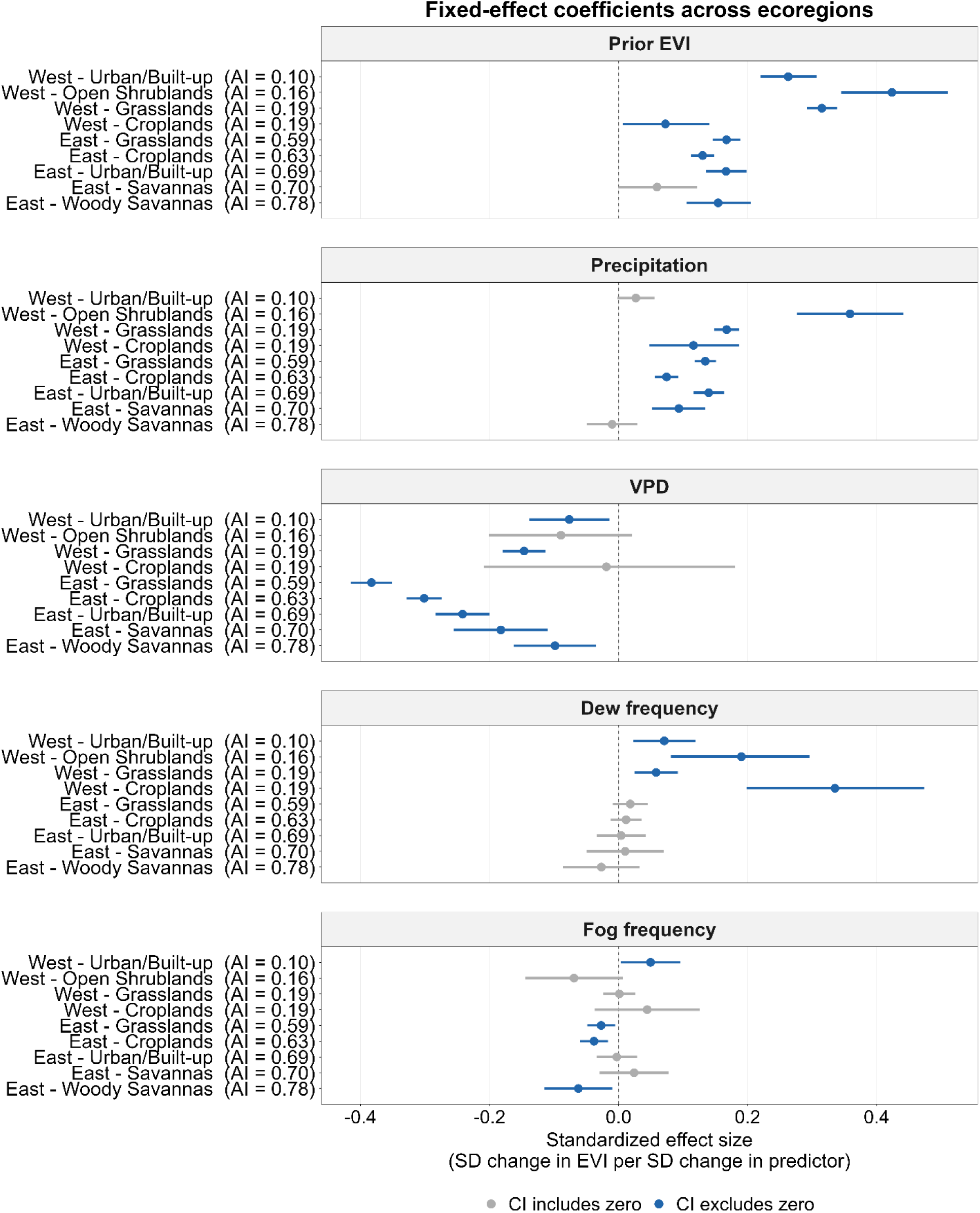
Fixed-effect coefficients from linear mixed-effects models predicting next-period EVI across nine U.S. ecoregions, ordered from most arid (top) to least arid (bottom). Points show standardized effect sizes; horizontal lines are 95% confidence intervals.

Increases in cumulative precipitation over a 16-day period were generally positively associated with vegetation growth, with this effect stronger in drier natural ecoregions and most pronounced in West Open Shrublands (*Fig. 3*). The only ecoregions where changes in precipitation did not have a significant effect on vegetation growth were East Woody Savannas (the most humid ecoregion) and West Urban/Built-up. Overall, we found precipitation coefficients to be fairly robust in all ecoregions (*Fig. S3*).

NRW had a less pronounced effect on vegetation compared to precipitation and VPD, although impacts of NRW were notable in many cases (*Fig. 3*). Specifically, increases in 16-day-period fog frequency had a clear positive effect in the West Urban/Built-up ecoregion, while negatively affecting vegetation in East Grasslands, East Croplands, and East Woody Savannas. As for dew frequency, it showed a clear positive effect to vegetation growth in all the water-limited western ecoregions, while its contribution was mostly neutral in the humid eastern ecoregions.

In our coefficient stability analysis the models behaved as expected when attributing the fog frequency coefficient, with mean VPD taking much but not all of the explanatory power away after adding the VPD variable to a fog-only model (*Fig. S4*). In West Open Shrublands, which hardly receive any fog during the summer, the fog coefficient was very unstable (*Fig. S4*), most likely due to the small sample size and outliers. For dew, likewise, mean VPD took much of the explanatory power away when added to a dew-only model (*Fig. S5*). Nevertheless, dew coefficient values remained always positive in the western ecoregions, regardless of the number of predictors included in the model (*Fig. S5*).

We lastly note that, even though dew frequency showed a neutral contribution in all the East ecoregions, it could be that dew is actually negatively impacting crops and other plants in humid ecoregions, but one of the meteorological conditions that contributes to dew formation, low cloud coverage, is having the opposite effect, thereby preventing us from observing the actual effect of dew. This hypothesis is supported by *Fig. S6*, where we refit the same model per biome as in *Fig. 3*, but this time also including mean hourly cloud coverage in oktas. In this case, all dew fixed-effect coefficients became slightly negative in the humid ecoregions, with 95% confidence intervals falling entirely below zero in East Croplands.

## 3. Discussion

### 3.1. Ecosystem-level water balance determines plant responses to NRW

Our study reveals widespread, non-negligible effects of summertime NRW on vegetation growth across the contiguous U.S., with plant responses primarily determined by the water availability of the system. By combining satellite-derived vegetation greenness data with weather-station observations, we expanded our ecosystem-scale understanding of how fog and dew affect vegetation. Such an ecosystem-level understanding is particularly needed for dew, since the often local, species-specific studies, the absence of publicly available dew datasets, and the high spatial heterogeneity of dew formation have hindered our ability to generalize on the effects of dew on ANPP across ecosystems and climatic gradients (2, 5).

Even after accounting for precipitation and VPD, we found that dew enhanced vegetation growth in water-limited systems of the West but was neutral or, sometimes even suppressed growth in humid systems of the East. Fog too had neutral or negative impacts in the more humid regions of our study, but, notably, its negative impacts were more widespread and intense than dew (*Figs. 2* and *3*). Importantly, even when models accounted for human modifications to vegetation that may occur within certain land-cover types, the aridity-dependent pattern of vegetation responses to NRW remained. That is, for a given land-cover type that exists both in the eastern and western U.S. (in our case, Urban/Built-up, Croplands, and Grasslands), dew and fog had a neutral-or-negative and a mostly positive effect on ANPP, respectively.

The above conclusions are further supported by the agreement of our results with the literature on how VPD and precipitation influence vegetation growth. For instance, our models showed that the influence of VPD on ANPP during the summer was considerable in humid climates, peaked in mesic climates, and then more sharply dropped and became negligible as aridity increased in semi-arid and arid locations. A similar pattern has been reported from eddy-covariance observational studies (12, 13), which show that humid and mesic ecosystems are largely constrained by VPD, with VPD limitations on growing-season productivity peaking in intermediately wet ecosystems, whereas soil-moisture limitations strengthen with increasing aridity and become dominant in semi-arid and arid regions. This pattern has been attributed to increasingly conservative water-use strategies in dry ecosystems, which reduce sensitivity to atmospheric demand and increase dependence on soil-water availability (13).

Our results on how summer precipitation influences ANPP also align well with prior ecological findings, which have shown that the sensitivity of vegetation to changes in precipitation increases toward drier and decreases toward wetter ecosystems (14, 15). Our models similarly showed that summer precipitation became increasingly important for vegetation growth when moving from humid to mesic and then semi-arid climates (*Fig 2*). Notably, however, the positive effects of dew on vegetation surpassed those associated with precipitation in the most arid sites of our study, where hardly any rainfall occurs over the dry summer season. This finding highlights the potential for NRW to maintain or even enhance plant health during dry spells.

### 3.2. Dew benefits vegetation more than fog in the water-limited West

In water-limited regions dew appears to have a more consistent benefit for vegetation than fog likely because, in the summer, dew occurs both along the coastline in higher frequencies and further inland in lower frequencies, whereas fog occurs almost exclusively along the coastline (*Fig. 1*). Physically, this is because summer fog forms over the cold Pacific Ocean along the West Coast under a strong marine inversion (16). As a result, fog rarely reaches far inland, and its influence in the West during the summer is limited to coastal ecosystems. In our study, fog showed a clear positive effect on vegetation in West Urban/Built-up vegetated areas, some of which are located along or near the West Coast (*Fig. 3*). This finding is consistent with field studies along the West Coast documenting benefits of summer advective fog on coastal ecosystems such as redwood forests (17, 18), pine forests (19), and coastal prairie grasslands (20).

In contrast, West Grassland and West Shrubland sites in our dataset are generally located far inland (*Fig. S1*), where summer fog is rare. For West Croplands specifically, results should be interpreted with caution, as our data included only 16 sites scattered across the western U.S. and mostly situated away from the coast. This limitation reflects the distribution of the ASOS weather stations in the Western U.S. As a result, fixed-effect estimates, although positive, were highly uncertain with wide confidence intervals that overlap zero (*Fig. 3*). We emphasize this because previous field studies have shown that summer fog can benefit crops in coastal California by increasing both water- and light-use efficiency (10, 21). Hence, it could be worthwhile for future efforts to further explore the large-scale impact of summer fog on crops by incorporating data from more agricultural fields along the California coast.

### 3.3. Negative impacts of fog and dew in the humid East

In the more humid areas encompassing much of the eastern U.S., we found that fog occurrence was widespread and that fog frequency had a more intense and extensive negative effect than dew in most ecoregions included in the study, negatively affecting ANPP in East Grasslands, East Croplands, and East Woody Savannas (*Fig. 3*). This observation could be due to three reasons. One reason is that fog prolongs leaf wetness, thereby creating favorable conditions for pathogen infection and increasing disease severity in already humid environments (22). Moreover, fog can scavenge atmospheric constituents and deposit them directly onto leaf surfaces, thus increasing surface water acidity and promoting nutrient leaching (23). Lastly, it is plausible that fog hinders plant growth in humid ecoregions due to decreasing the number of hours with sunshine in areas that already receive frequent precipitation and cloud coverage and are therefore less water- and more light-limited (24).

The reason why dew was initially not found to negatively affect plant growth in humid (especially agricultural) ecoregions at the spatiotemporal scale of our analysis is not entirely clear, as dew also prolongs leaf wetness, thereby promoting pathogen growth, and can deposit atmospheric constituents onto leaves, thereby promoting nutrient leaching. The most plausible explanation is the indirect offset of the negative effects by a factor that immediately affects dew: cloud coverage. More specifically, cloudless hours around dawn provide favorable conditions for dew formation, as radiative cooling of the Earth’s surface is high (11). Since the equation used in the present study to estimate dew had cloud coverage as one of its inputs, we hypothesize and found evidence (*Fig. S6*) that it was cloudless conditions that indirectly offset the negative effects of dew from being observed in some humid ecoregions, as cloudless post-dawn hours (especially when water is not limiting) would increase plant photosynthetic activity (24).

Our findings of negative impacts of NRW on ANPP in East Cropland sites (*Fig. S6*), combined with observations of increasing summer dewpoint over the past 70 years in much of the Midwest (25), suggest that emphasis should be placed on monitoring summertime NRW in eastern U.S. croplands. Forecasting, monitoring trends, and developing strategies to reduce the frequency or persistence of dew and fog could help mitigate crop yield losses in one of the most productive agricultural regions of the U.S. Conversely, in water-limited systems of the western U.S., where intensifying droughts and water scarcity increasingly threaten irrigated agriculture (26–29), efforts could focus on maximizing the beneficial effects of dew and fog during the summer. This could be achieved either by the use of artificial dew after identifying the specific species that benefit from it, or by breeding crops that better utilize NRW.

### 3.4. Fog and dew deserve more attention in ecohydrology

Overall, our study provides the first evidence of noteworthy effects of summertime NRW on vegetation growth at the ecosystem level and over relatively long (that is, monthly) time spans. Given our observed differential impacts of summer fog and dew on vegetation, a promising future direction would be to expand the study of NRW dynamics and ecosystem-level impacts across all four seasons. More broadly, we conclude that our understanding of ecohydrology would become more complete if more ecological and agricultural studies were to begin incorporating this much-overlooked component of the hydrologic cycle. Furthermore, because it remains largely unclear how climate change is affecting fog and dew across the U.S., future efforts aimed at improving our understanding of these changes could also provide a more complete picture of vegetation responses amid increasing hydroclimate volatility and water scarcity (30, 31).

## 4. Materials and Methods

### 4.1. Overview

We combined satellite-derived vegetation greenness (MODIS EVI) with ASOS weather station meteorological data to assess the role of summertime NRW (specifically fog and dew) across U.S. regions of varying water availability. Changes in vegetation greenness across 16-day summer periods were modeled by regressing EVI at period *t+1* on lagged EVI at period *t−1*, together with precipitation amount, mean vapor pressure deficit, fog frequency, dew frequency, and mean cloud coverage at period *t*, and a station-level random intercept. This linear mixed-effects modeling approach was applied continuously along the aridity gradient using 30 equal-sized station bins (36 stations per bin), and resulting coefficients were related to aridity using generalized additive models. The approach was also applied categorically across distinct ecoregions characterized by land-cover class and an eastern–western split at the 100th meridian. To assess the robustness of inferred fixed effects within each ecoregion, we conducted a coefficient stability analysis by fitting all possible subsets of hydroclimatic predictors and evaluating the variability of fixed-effect estimates across models.

### 4.2. Data Sources

Hourly U.S. weather station data for the years 2012–2023 were obtained from the Automated Surface Observing System (ASOS) network and accessed via the Iowa Environmental Mesonet (IEM) archive (32). These data included hourly air temperature (converted to °C), dew point temperature (converted to °C), precipitation (converted to mm), wind speed (converted to m/s), cloud coverage (converted to oktas), and fog/mist occurrence flags. Vapor pressure deficit (VPD) was calculated at the hourly scale from air and dew point temperatures using the Tetens formulation (33).

Vegetation data for ∼500-m pixels containing each ASOS station were obtained from the Moderate Resolution Imaging Spectroradiometer (MODIS) Enhanced Vegetation Index (EVI) 16-day composite product (MOD13A1 Version 6.1) (34). Land cover class (∼500-m resolution) was assigned from the MODIS MCD12Q1 IGBP product (35).

Monthly aridity index (AI) data for ∼1000-m pixels containing each station were obtained from the Global Aridity Index and Potential Evapotranspiration (ET0) Database Version 3.1, representing a water-balance climatology for 1970-2000 (36). The AI is defined as the ratio of monthly precipitation to monthly potential evapotranspiration. An average monthly summer AI value was derived for each station by averaging June, July, and August monthly values, with lower values indicating more arid conditions.

### 4.3. Data Processing

Hourly ASOS observations were first filtered to remove erroneous hourly temperature and daily precipitation values exceeding state-level historical extremes derived from NOAA NCEI State Climate Extremes records (37). Climatic data were then aggregated into five MODIS 16-day summer periods (days of year 161–176, 177–192, 193–208, 209–224, and 225–240). Within each period, precipitation was summarized as total amount (mm) and frequency (percentage of hours with ≥0.25 mm). Fog and mist occurrence were expressed as fog frequency (percentage of hours with fog/mist), while cloud coverage, air temperature and VPD were averaged in each period across all valid hourly observations. Station-years were retained only if all five summer periods within a given year met a 90% hourly data completeness threshold, and only stations with at least 8 years of complete summer coverage during 2012–2023 were included in the analysis.

Dew formation was estimated from hourly local observations as potential dew yield using an analytical formula based on radiative cooling and atmospheric conditions, which included air temperature, dew point temperature, wind speed, cloud cover, and elevation (11). An hour’s dew was set to zero if precipitation (> 0.25 mm) or fog occurred during that hour. Dew was then aggregated within each 16-day period as estimated total dew amount (mm) and estimated dew frequency (percentage of hours with dew occurrence). We note here that, due to differences in the surface properties of the aboveground tissues of different plant species and differences in the microenvironmental conditions in the atmosphere immediately surrounding the canopy, the equation herewith used is not expected to provide perfectly accurate dew measurements. Rather, this equation identifies hours at a station when dew is very likely to occur in its vicinity.

MODIS EVI observations were filtered using quality assurance flags to retain only good- or marginal-quality pixels with acceptable usefulness scores and to exclude observations affected by clouds and other artifacts. Subsequently, stations where 50% or more of their MODIS periods fell below the EVI threshold of 0.1 were excluded from the analysis, as we assumed they lacked sufficient surrounding vegetation. Also, even for stations that passed this filtering criterion, only periods with EVI ≥ 0.1 were retained. This was done to exclude periods when extreme disturbances (e.g., harvesting) in managed landscapes would substantially alter EVI.

For the ecoregion modeling only (section 4.5), stations where the MODIS land-cover class was not consistent across all years of the study period were excluded, as interannual reclassification of a station to a different land-cover class would introduce ambiguity in ecoregion assignment.

### 4.4. NRW Effects Along the Aridity Gradient

To examine how the effects of precipitation amount, mean VPD, dew frequency, and fog frequency on vegetation vary along the summer aridity gradient, the 1,080 stations that passed the filtering criteria were ranked by their summer AI value and divided into 30 equal-sized bins of 36 stations each. Within each bin, all predictors and the response variable were standardized (z-scored) within each 16-day period to account for systematic differences in mean climatic conditions and vegetation greenness across the 5 summer periods. A linear mixed-effects model was then fit separately for each bin. The model took the form:

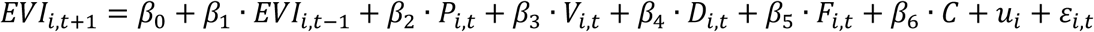

where *i* indexes stations and *t* indexes 16-day periods; *P* is precipitation amount, *V* is mean VPD, *D* is dew frequency, *F* is fog frequency, and *C* is average cloud coverage; *u_i_*∼*N*(0, *σ*^2^) is a station-level random intercept; and *ε_i_*_,*t*_∼*N*(0, *σ*^2^) is the residual error.

We decided to exclude mean air temperature, precipitation frequency, and dew amount from the modeling process because they were highly correlated with mean VPD, precipitation amount, and dew frequency, respectively (Pearson correlation coefficients between 0.62– 0.94, 0.50–0.95, and 0.88–0.96, respectively, depending on the bin).

Following the modeling in each bin, fixed-effect coefficients were extracted and mapped spatially across the contiguous U.S. A generalized additive model (GAM) was then fit separately for each hydrological variable, with bin-level fixed-effect estimates as the response and the bin mean summer AI as the predictor, using a thin plate regression spline (k = 5) with the smoothing parameter selected automatically by REML and estimates weighted by inverse variance to give greater influence to bins with more precise estimates. For each GAM, using an approximate test of the smooth term we assessed whether the fitted curve differed significantly from a flat line at zero, such that a small p-value indicates a systematic relationship between the coefficient (i.e., the effects of a hydrological variable on vegetation growth) and aridity. Conducting the same analysis across bin sizes ranging from 20 to 40 (27–54 stations per bin) confirmed qualitative robustness of results. Hence, results from 30 bins are reported throughout this paper.

As a final quality-control step, we applied a leave-one-station-out influence analysis to each bin’s model, where we refit the same model with one station removed at a time and calculated the standardized shift in each fixed-effect estimate (DFBETAS) to quantify that station’s influence. Every station exceeding a DFBETAS magnitude of 1 for a given predictor (i.e., removing that station shifted the coefficient by more than one standard error) was manually checked against its 16-day-period meteorological record. In this way, we identified one station (Harvey, ND) with an implausible precipitation value (3,218.69 mm within a single 16-day period) repeated identically across four periods, followed by several years of anomalous zero-precipitation reporting. This station was excluded from the analysis. All other flagged stations showed no corresponding anomaly and were retained.

Although we do expect some of the weather stations to contain surrounding vegetation that is managed (e.g., through irrigation or mowing), we do nonetheless expect that not all anthropogenic disturbances will be synchronized between stations, allowing the large sample size within each bin to robustly estimate the effects of the hydrological variables on vegetation growth. Moreover, any persistent station-level differences in vegetation productivity not explained by the hydrological predictors (such as those arising from soil properties, topography, or constant irrigation) are absorbed by the station-level random intercept, preventing them from confounding the fixed-effect estimates. Lastly, prior-period EVI that is included in the modeling should account at least to some extent for vegetation memory.

### 4.5. Ecoregion Modeling

To assess whether the observed, aridity-dependent effects of fog and dew on vegetation growth persisted after accounting for land-cover type, as well as to make these effects easier to interpret across well-defined and widely recognized ecoregions, 856 stations were partitioned into ecoregions based on land-cover type and a western-eastern split using the 100^th^ meridian—a well-established boundary separating water-limited and humid climates in the U.S. (38). Separate linear mixed-effects models were fit for each ecoregion, with all response and predictor variables standardized within each ecoregion and 16-day period. The model took the form:

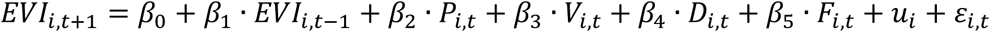

where *i* indexes stations and *t* indexes 16-day periods; *P* is precipitation amount, *V* is mean VPD, *D* is dew frequency, and *F* is fog frequency; *u_i_*∼*N*(0, *σ*^2^) is a station-level random intercept; and *ε_i_*_,*t*_∼*N*(0, *σ*^2^) is the residual error.

Models were fit only to ecoregions with at least 10 stations. We decided to exclude mean air temperature, precipitation frequency, and dew amount from the modeling process because they were highly correlated with mean VPD, precipitation amount, and dew frequency, respectively (Pearson correlation coefficients between 0.62 – 0.90, 0.61 – 0.79, and 0.91 – 0.97, respectively, depending on the ecoregion).

### 4.6. Coefficient Stability Analysis

To evaluate the robustness of the fixed-effect coefficients for the models of each ecoregion, we conducted a coefficient stability analysis. This was particularly done to identify cases in which a model could not precisely attribute the independent contributions of VPD and fog, or VPD and dew, to plant growth, given that these predictors can be somewhat correlated.

Within each ecoregion, we fit mixed-effects linear models for all possible subsets of hydrological predictors while retaining lagged EVI and the station-level random effect. This resulted in 15 model variants per ecoregion, with each predictor included in 8 of these models. For each hydroclimatic predictor in each ecoregion, we extracted fixed-effect estimates and assessed their stability by examining the distribution of coefficient values across all model variants that included that predictor.

## Data, Materials, and Software Availability

ASOS weather observations are publicly available through the Iowa Environmental Mesonet archive. MODIS EVI and land-cover products are publicly available through NASA LP DAAC. The Quarto workflow and associated R code used for statistical analyses and figure generation, together with the processed datasets required to reproduce the analyses, are publicly available on the Open Science Framework (OSF) at https://doi.org/10.17605/OSF.IO/K5JFQ. All code was written and executed in R version 4.5.1 (39).

## Acknowledgements

The authors acknowledge Research Computing at Arizona State University for providing High-Performance Computing (HPC) resources that have contributed to the research results reported within this paper (40).

