## Supplemental Figures and Tables for "Summertime Fog and Dew Impacts on Vegetation Depend on Aridity"

**File includes:**

Figs. S1 to S6

Tables S1 and S2

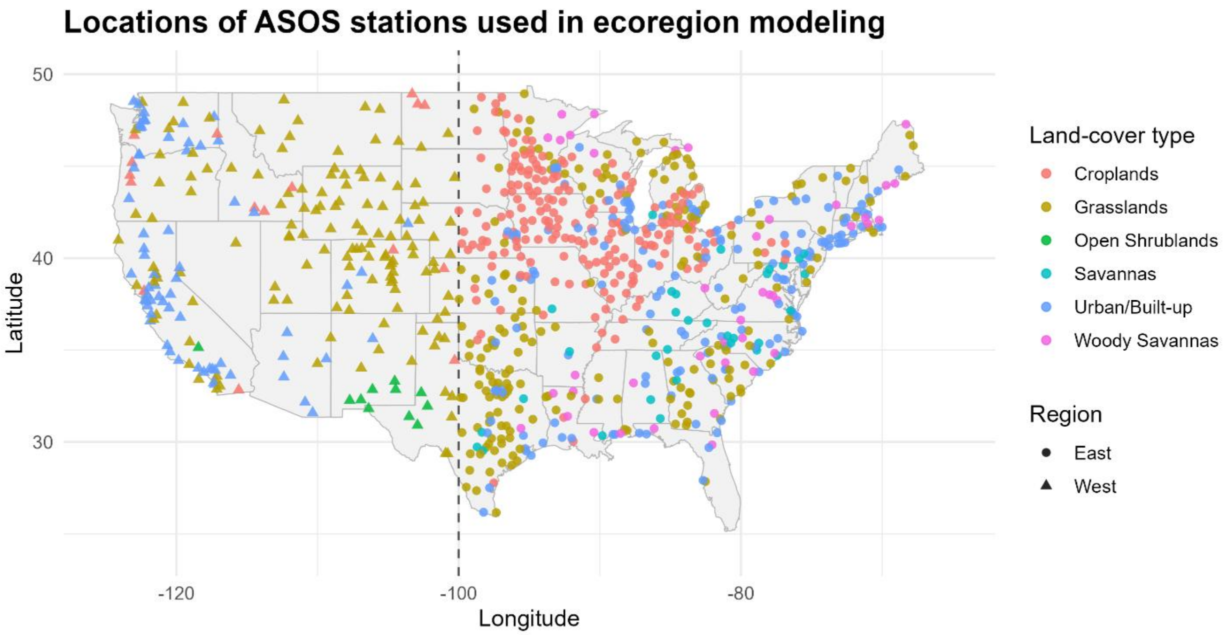


***Fig. S1.*** *Locations of the 856 ASOS weather stations included in the ecoregion modeling process after all filtering criteria were applied.*

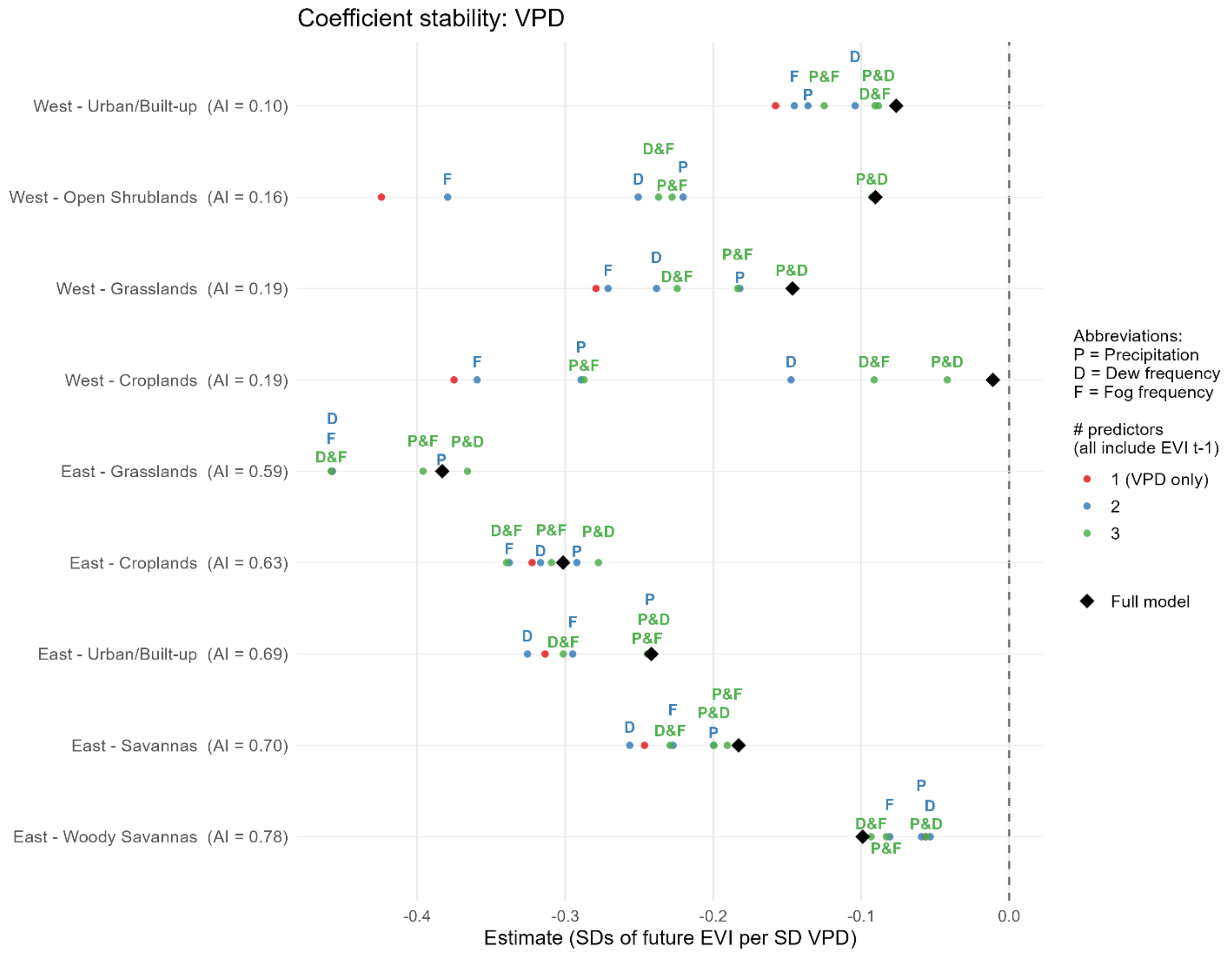


***Fig. S2.*** *Coefficient stability analysis for mean VPD across ecoregions. Each point represents the estimated fixed effect from one of 15 mixed-effects model variants, each including* $EVI_{(t-1)}$ *and a station-level random intercept but differing in which subset of the four predictors is included. Red dots represent univariate models in which mean VPD appears alongside EVI(t−1) only, with no other predictors controlled for. Blue and green dots represent 2- and 3-predictor models respectively, with letters identifying the other predictors included (P = Precipitation Amount, D = Dew Frequency, F = Fog Frequency). The black rhombus marks the full model estimate containing all four predictors. Ecoregions are ordered from most arid (top) to most humid (bottom) by mean summer aridity index.*

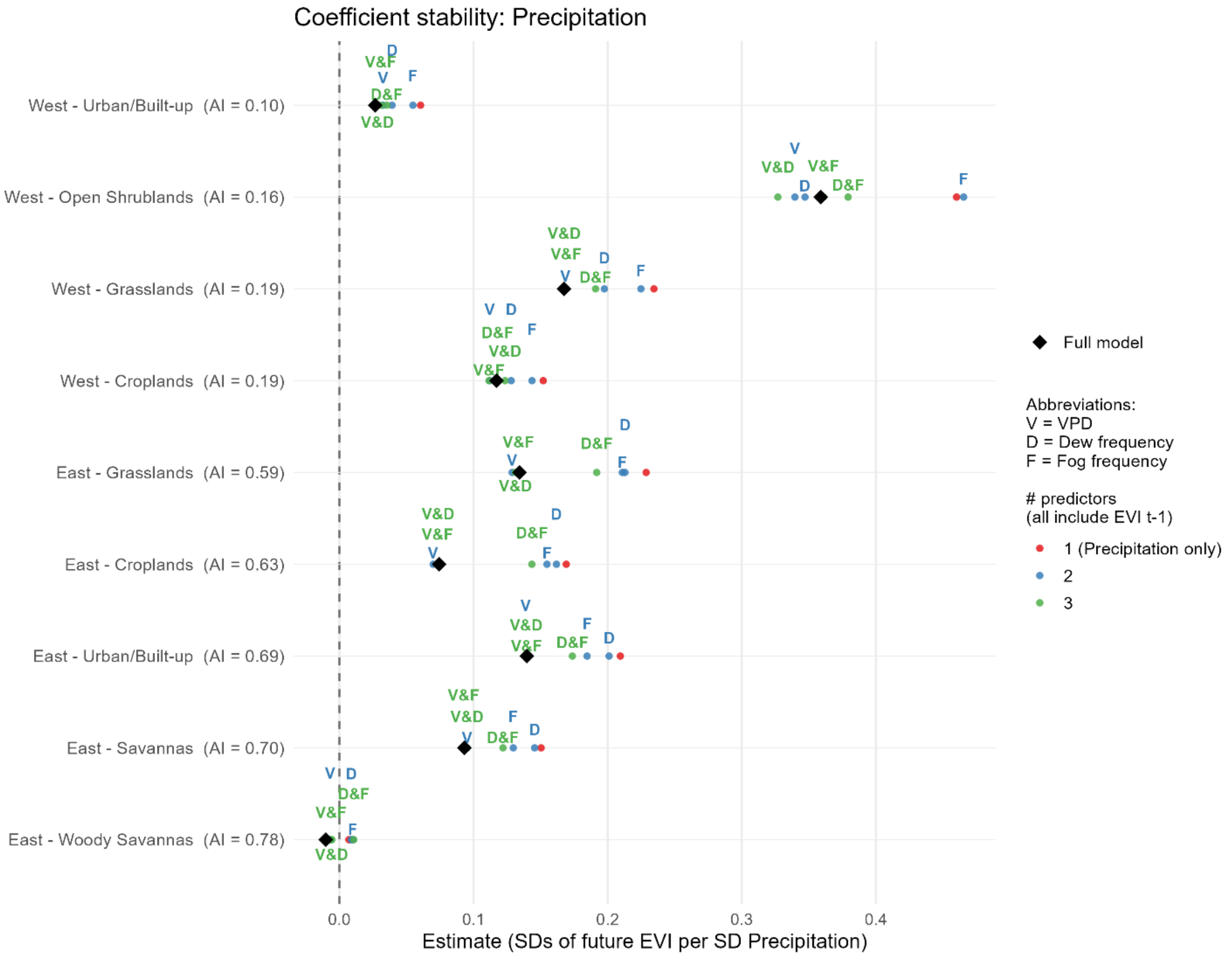


***Fig. S3.*** *Coefficient stability analysis for total precipitation across ecoregions. Each point represents the estimated fixed effect from one of 15 mixed-effects model variants, each including* $EVI_{(t-1)}$ *and a station-level random intercept but differing in which subset of the four predictors is included. Red dots represent univariate models in which precipitation appears alongside EVI(t−1) only, with no other predictors controlled for. Blue and green dots represent 2- and 3-predictor models respectively, with letters identifying the other predictors included (V = VPD, D = Dew Frequency, F = Fog Frequency). The black rhombus marks the full model estimate containing all four predictors. Ecoregions are ordered from most arid (top) to most humid (bottom) by mean summer aridity index.*

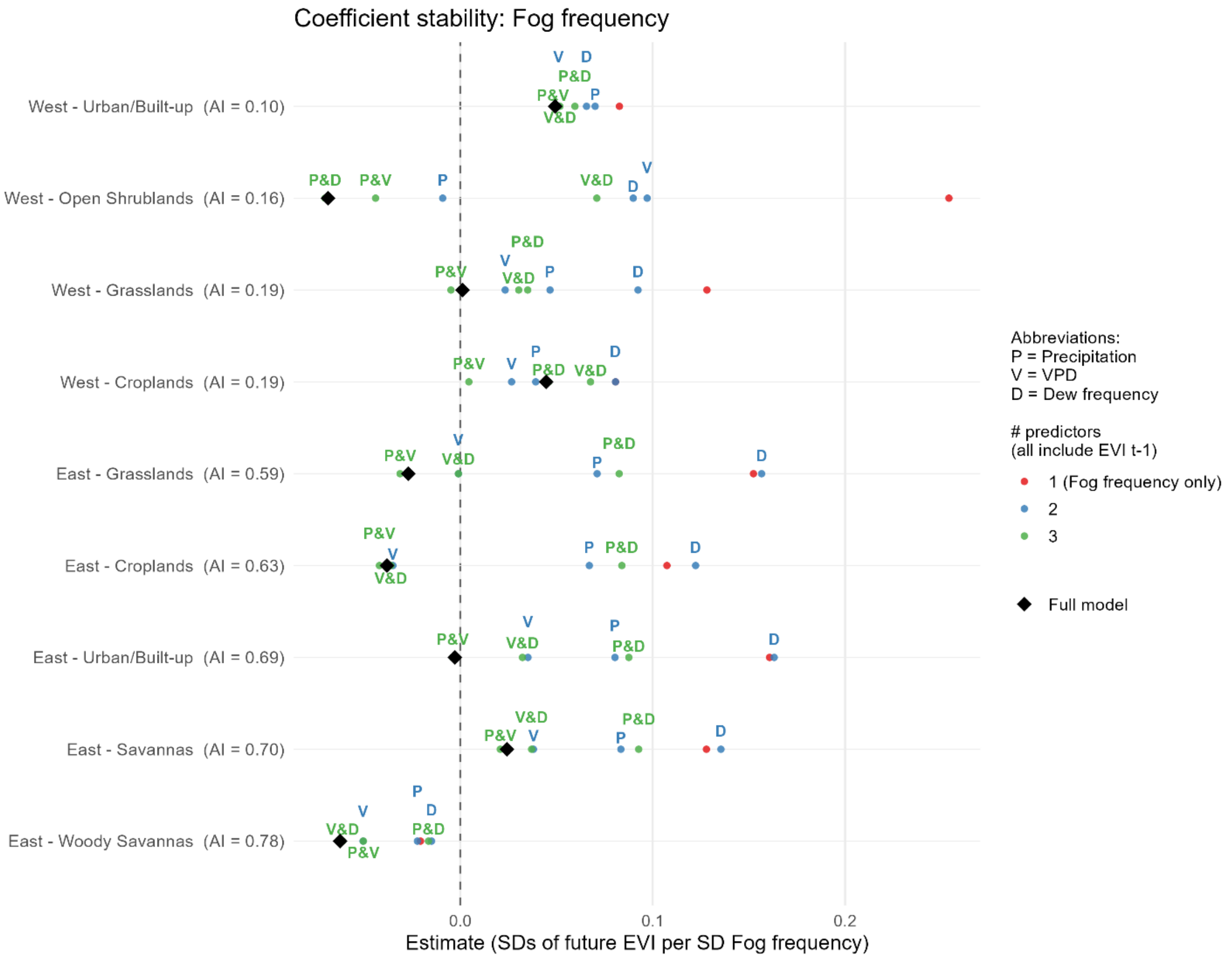


***Fig. S4.*** *Coefficient stability analysis for fog frequency across ecoregions. Each point represents the estimated fixed effect from one of 15 mixed-effects model variants, each including* $EVI_{(t-1)}$ *and a station-level random intercept but differing in which subset of the four predictors is included. Red dots represent univariate models in which fog frequency appears alongside EVI(t−1) only, with no other predictors controlled for. Blue and green dots represent 2- and 3-predictor models respectively, with letters identifying the other predictors included (P = Precipitation Amount, V = VPD, D = Dew Frequency). The black rhombus marks the full model estimate containing all four predictors. Ecoregions are ordered from most arid (top) to most humid (bottom) by mean summer aridity index.*

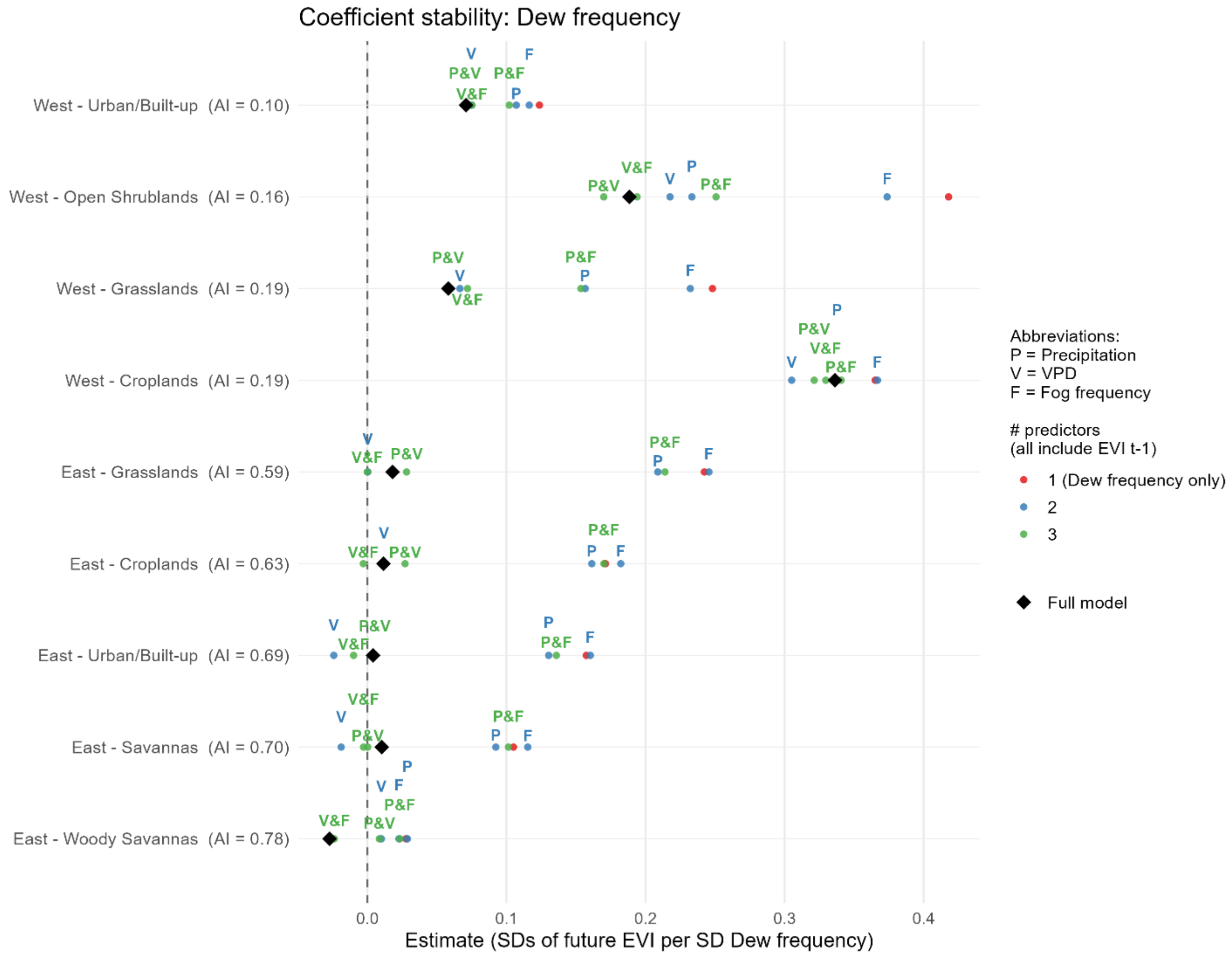


***Fig. S5.*** *Coefficient stability analysis for dew frequency across ecoregions. Each point represents the estimated fixed effect from one of 15 mixed-effects model variants, each including* $EVI_{(t-1)}$ *and a station-level random intercept but differing in which subset of the four predictors is included. Red dots represent univariate models in which dew frequency appears alongside EVI(t−1) only, with no other predictors controlled for. Blue and green dots represent 2- and 3-predictor models respectively, with letters identifying the other predictors included (P = Precipitation Amount, V = VPD, F = Fog Frequency). The black rhombus marks the full model estimate containing all four predictors. Ecoregions are ordered from most arid (top) to most humid (bottom) by mean summer aridity index.*

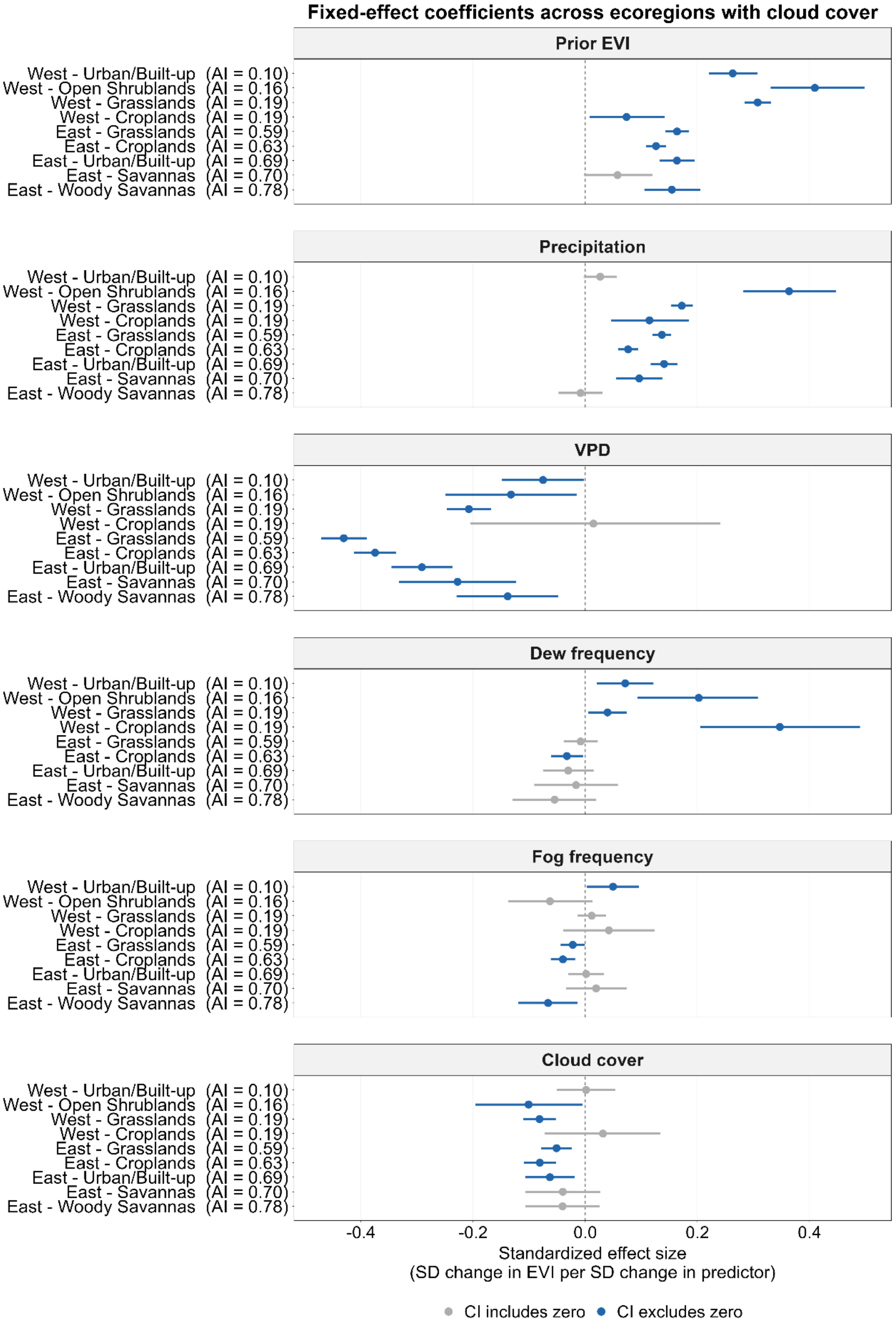


***Fig. S6.*** *Models fitted exactly as in Figure 3, but this time also including average hourly cloud coverage (in oktas) within a 16-day period.*

| Aridity bin | Mean Summer AI | $R^2$ marginal | $R^2$ conditional |
|---|---|---|---|
| 1 | 0.01 | 0.27 | 0.48 |
| 2 | 0.04 | 0.04 | 0.61 |
| 3 | 0.11 | 0.05 | 0.62 |
| 4 | 0.17 | 0.30 | 0.60 |
| 5 | 0.21 | 0.10 | 0.60 |
| 6 | 0.25 | 0.39 | 0.70 |
| 7 | 0.29 | 0.43 | 0.55 |
| 8 | 0.31 | 0.30 | 0.58 |
| 9 | 0.35 | 0.30 | 0.53 |
| 10 | 0.41 | 0.28 | 0.61 |
| 11 | 0.47 | 0.18 | 0.58 |
| 12 | 0.55 | 0.11 | 0.51 |
| 13 | 0.58 | 0.10 | 0.60 |
| 14 | 0.60 | 0.11 | 0.59 |
| 15 | 0.61 | 0.22 | 0.56 |
| 16 | 0.63 | 0.07 | 0.64 |
| 17 | 0.65 | 0.12 | 0.58 |
| 18 | 0.66 | 0.16 | 0.60 |
| 19 | 0.67 | 0.10 | 0.55 |
| 20 | 0.69 | 0.14 | 0.55 |
| 21 | 0.70 | 0.13 | 0.67 |
| 22 | 0.71 | 0.09 | 0.61 |
| 23 | 0.72 | 0.13 | 0.56 |
| 24 | 0.74 | 0.14 | 0.55 |

| | | | |
|---|---|---|---|
| 25 | 0.76 | 0.12 | 0.57 |
| 26 | 0.78 | 0.13 | 0.52 |
| 27 | 0.81 | 0.06 | 0.68 |
| 28 | 0.84 | 0.10 | 0.54 |
| 29 | 0.92 | 0.06 | 0.59 |
| 30 | 1.09 | 0.08 | 0.59 |

***Table S1.*** *Model performance by aridity bin.*

| Ecoregion | Station-sites (n) | Station-period observations (N) | $R^2$ marginal | $R^2$ conditional |
|---|---|---|---|---|
| East Croplands | 200 | 8,654 | 0.15 | 0.44 |
| East Grasslands | 196 | 7,149 | 0.33 | 0.59 |
| East Savannas | 34 | 1,049 | 0.10 | 0.63 |
| East Urban/Built-up | 158 | 3,941 | 0.17 | 0.52 |
| East Woody Savannas | 41 | 1,448 | 0.04 | 0.56 |
| West Croplands | 16 | 707 | 0.15 | 0.47 |
| West Grasslands | 134 | 5,695 | 0.31 | 0.60 |
| West Open Shrublands | 11 | 403 | 0.51 | 0.60 |
| West Urban/Built-up | 66 | 2,125 | 0.17 | 0.59 |

***Table S2.*** *Model performance and sample size by ecoregion.*
